# Engineering the thumb loop of T7 RNA polymerase improves the integrity of long RNA transcripts

**DOI:** 10.64898/2026.11.10.765520

**Authors:** Wei He, Hongli Zhou, Guiying Ji, Xiaoyu Xu, Qiuheng Jin

## Abstract

Incomplete transcripts complicate therapeutic mRNA manufacturing, particularly for long constructs such as self-amplifying RNA (saRNA). Their mechanisms remain poorly understood, hindering targeted improvements to mRNA integrity, a critical quality attribute. Here, our goal was to engineer the thumb loop of T7 RNA polymerase (RNAP) to improve transcript integrity. Using templates derived from a common 12 kb RNA sequence, we observed decreasing integrity as transcript length increased. Substitution of proline residues within the thumb loop improved integrity across 4, 8, and 12 kb transcripts, with several variants outperforming the previously reported K389A variant on the 12 kb template. Further substitutions identified M362K as the top performer and K363R as beneficial, implicating local conformation and electrostatic interactions. M362K retained its integrity-enhancing effect with N^1^-methylpseudouridine or 5-methylcytidine incorporated during transcription. These findings identify the thumb loop as a target for improving the integrity of long, chemically modified RNA transcripts.

---

mRNA applications are expanding from prophylactic vaccines to cancer immunotherapy, protein replacement, and in vivo cell and gene engineering.^1^ This broad therapeutic scope places increasing demands on manufacturing processes that consistently deliver high-quality RNA. However, mechanistic understanding and targeted control of transcription-related impurities have advanced unevenly. For double-stranded RNA (dsRNA), studies have characterized its formation pathways and linked them to specific enzymatic activities.^2, 3^ These insights have informed polymerase engineering^4, 5^ and targeted process interventions, including chaotropic additives^6^ and site-specific non-template DNA nicking,^7^ to suppress dsRNA production. Despite constituting the largest RNA impurity fraction, incomplete transcripts remain mechanistically poorly understood. This knowledge gap hinders targeted efforts to improve mRNA integrity, a critical quality attribute.

Characterization of incomplete BNT162b2 transcripts revealed preserved 5’ cap structures but absent poly(A) tails, implicating premature transcription termination as their predominant source.^8^ However, these findings do not explain the molecular events that prevent transcriptional completion. This question becomes increasingly relevant as therapeutic applications may require mRNAs encoding large proteins, such as Cas9. Additional regulatory elements, including replicase modules of saRNA, can further extend transcript length.^9^ Longer transcripts demand sustained polymerase processivity over greater distances, potentially increasing the likelihood of premature termination and the burden of incomplete products. These challenges motivate strategies to improve transcriptional completion during RNA synthesis.

To examine the relationship between transcript length and integrity, we designed a panel of DNA templates encoding a 12-kb RNA transcript and shorter segments of the same RNA sequence. These included the 5’-terminal 8- and 6-kb segments, three contiguous, non-overlapping 4-kb segments, and six contiguous, non-overlapping 2-kb segments (Fig. 1A). Each set of equal-length segments collectively spanned the entire 12-kb RNA transcript. Among transcripts sharing the same 5’ end, integrity progressively decreased as transcript length increased from 2 to 12 kb, indicating a negative association between length and integrity (Fig. 1B). No statistically significant differences in integrity were detected among the six 2 kb transcripts or among the three 4-kb transcripts (Fig. 1C, D). Together, these results support transcript length as an important determinant of integrity within this template panel.

**Figure 1.**
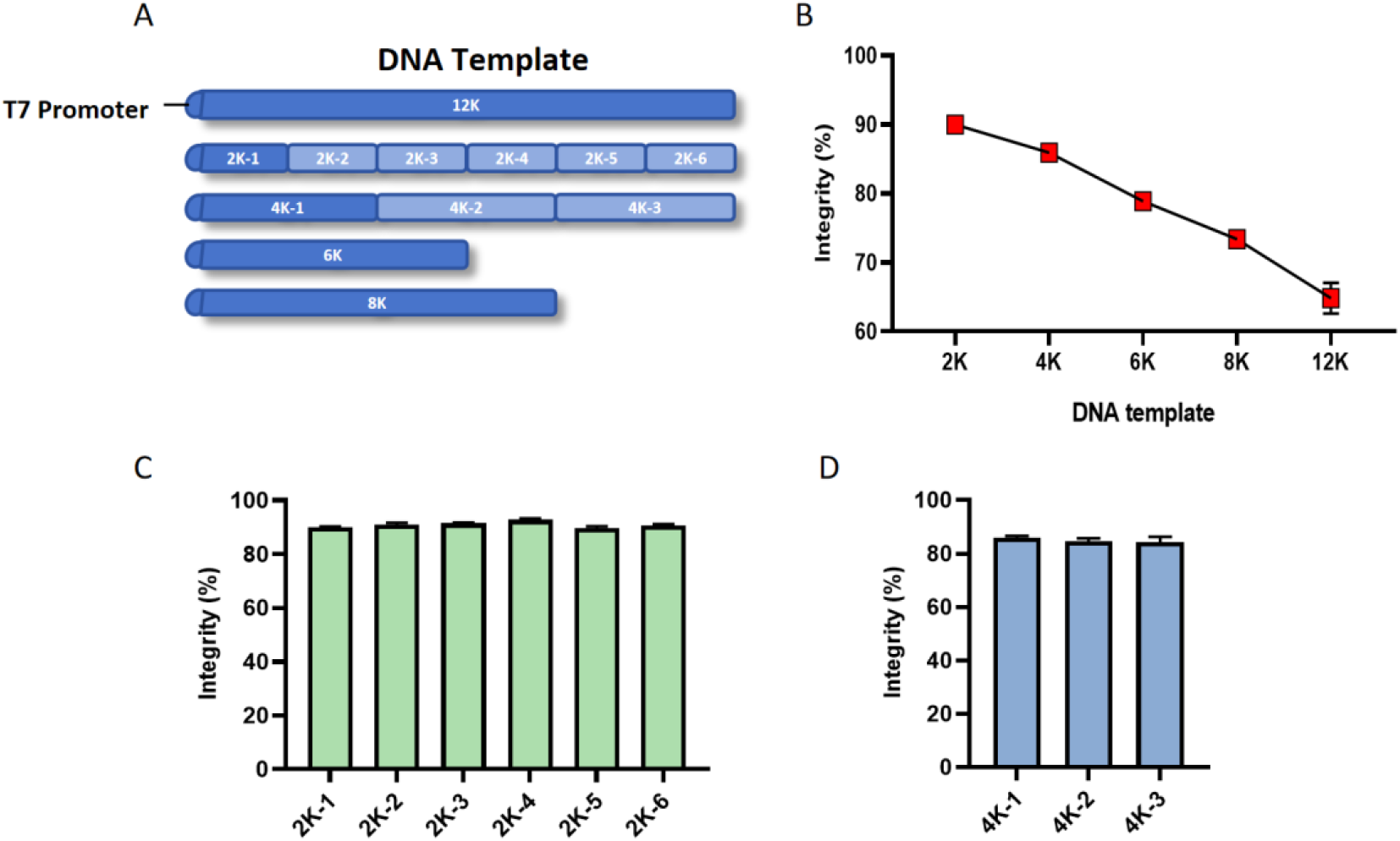
Measurement of RNA integrity. (A) Schematic of the DNA template; (B) RNA product integrity decreases as DNA template increases; (C) Measurement of RNA product integrity using 2-kb DNA templates; (D) Measurement of RNA product integrity using 4-kb DNA templates.

The length-dependent decline in integrity, together with evidence implicating premature termination in incomplete transcript formation, prompted us to ask whether maintaining elongation-complex (EC) stability could improve transcriptional completion. Longer transcripts require more nucleotide-addition cycles, potentially increasing the cumulative opportunity for premature termination. Structural studies of T7 RNAP show that the RNA–DNA hybrid is enclosed within a protein cavity and that rearrangement of the N-terminal domain establishes an RNA exit pathway.^10^ Biochemical studies further implicate the thumb subdomain and its interactions with the fingers subdomain in EC stability and termination.^11, 12^ Consistent with the thumb subdomain being a candidate target for engineering, K389A, a substitution in its helix N, was previously reported to improve mRNA integrity.^13^

We identified a relatively long loop containing multiple prolines, spanning residues 343–373 within the thumb subdomain (Fig. 2A). Because proline constrains local backbone conformations, we substituted the proline residues in this region with alanine or leucine to test whether changes in local conformation could improve transcript integrity. We tested these variants on the 4K-1, 8K, and 12K templates. Most variants improved integrity, with greater improvements observed for longer transcripts (Fig. 2B-D). On the 12K template, several loop variants produced substantially greater improvements than K389A. This previously reported integrity-enhancing variant provided only a modest benefit under the same conditions (Fig. 2D). These results identify the thumb loop as a promising target for improving the integrity of long RNA transcripts.

**Figure 2.**
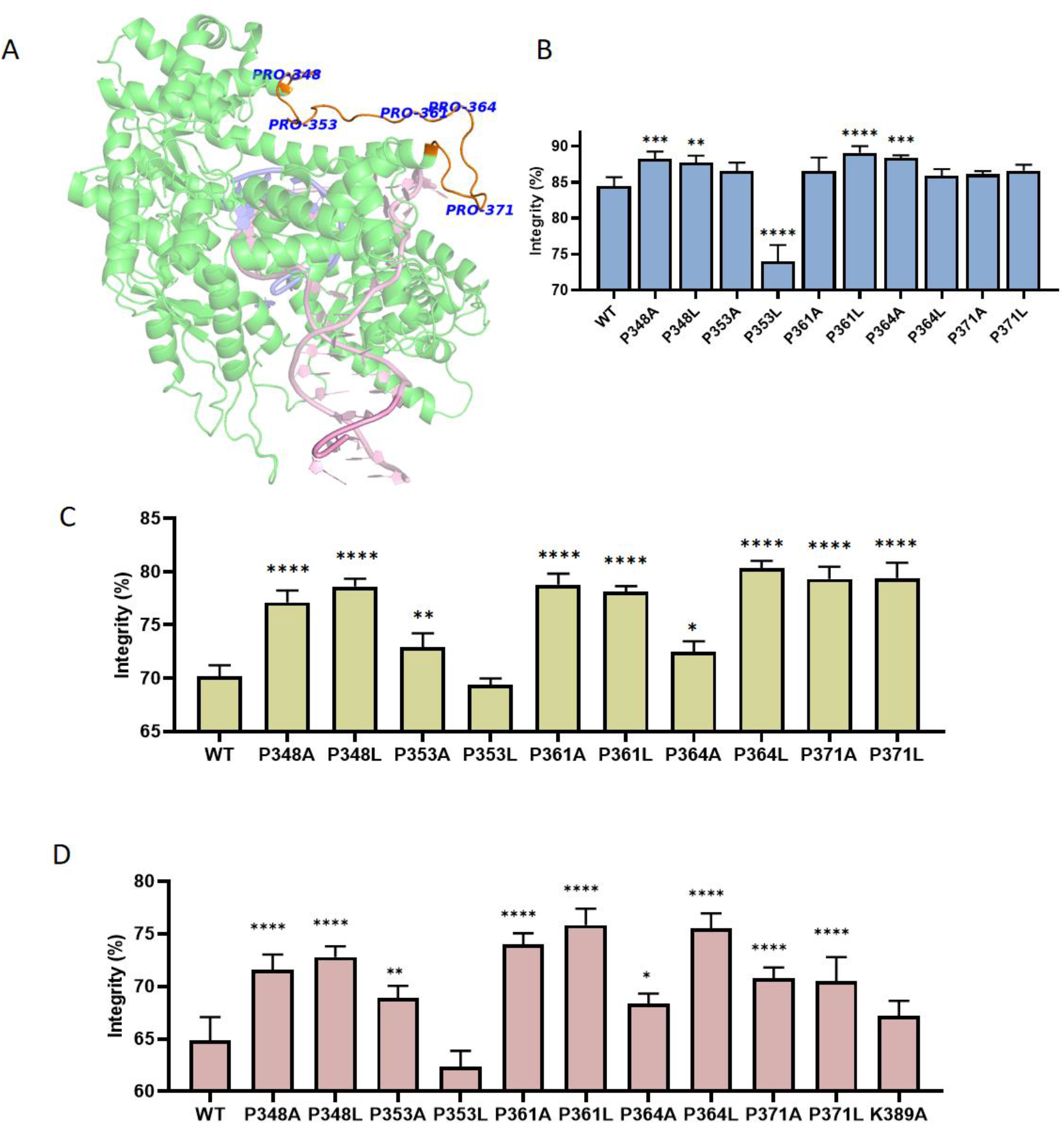
Substitutions improved integrity of RNA products. (A) Schematic diagram showing the positions of proline residues in the loop; (B) Measurement of RNA product integrity using a 4-kb DNA template; (C) Measurement of RNA product integrity using an 8-kb DNA template; (D) Measurement of RNA product integrity using a12-kb DNA template.

Substitutions at P361 and P364 improved integrity more than those at the other tested proline positions, directing our attention to the intervening residues M362 and K363. Mapping this region onto the T7 RNAP structure placed it near the upstream DNA exit region (Fig. 3A). We introduced positively charged residues at M362 and replaced K363 with alanine or arginine. M362K, M362R, and K363R significantly improved integrity, among which M362K showed the greatest improvement, exceeding that obtained with the tested proline substitutions, whereas K363A did not reduce it (Fig. 3B, C). These results suggest that local electrostatic properties contribute to the function of this region.

**Figure 3.**
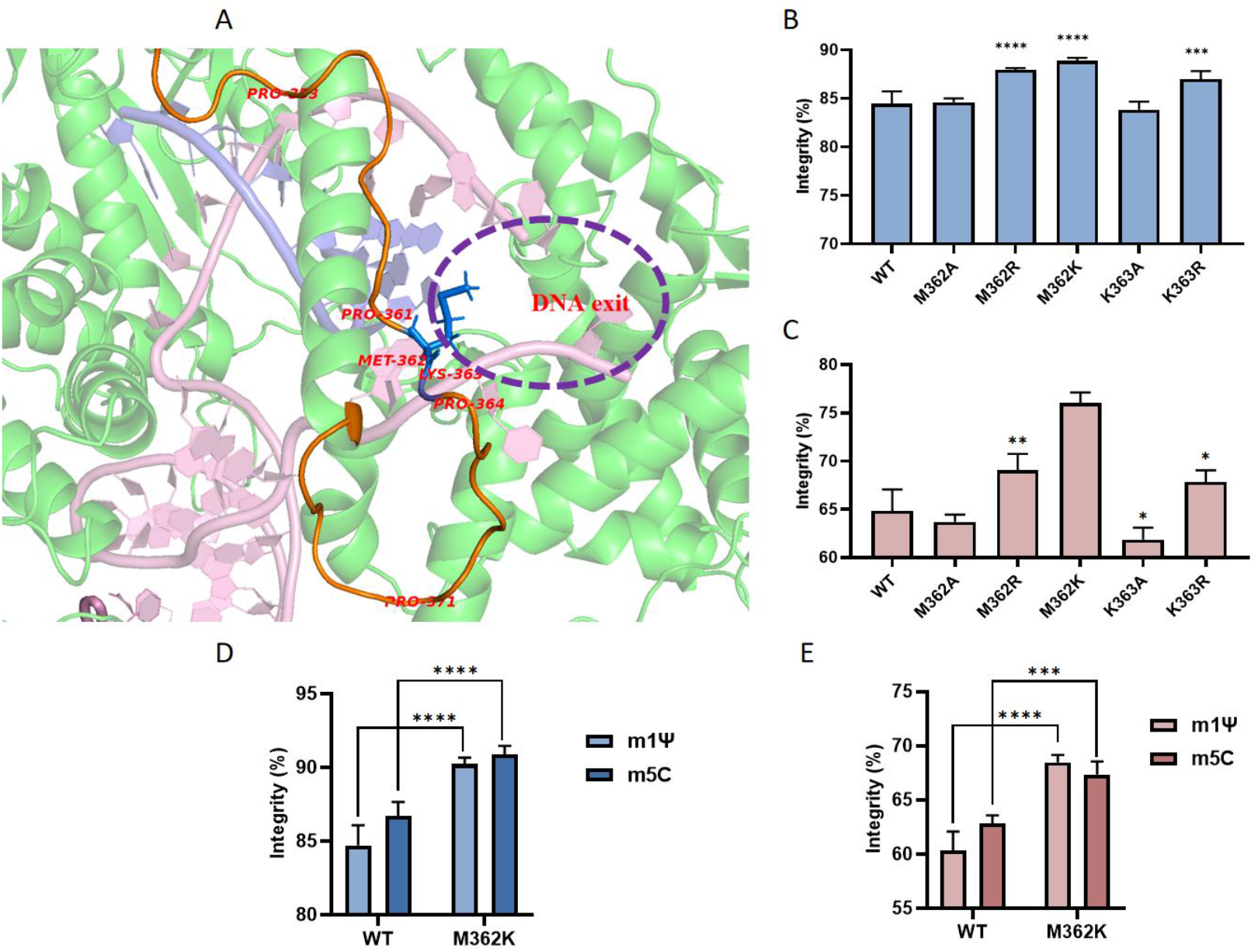
Substitutions further improve RNA product integrity. M362–K363 near the T7 RNAP upstream DNA exit; (B) Measurement of RNA product integrity using a 4-kb DNA template with unmodified nucleotides as substrates; (C) Measurement of RNA product integrity using a 12-kb DNA template with unmodified nucleotides as substrates; (D) Measurement of RNA product integrity using a 4-kb DNA template with modified nucleotides as substrates; (E) Measurement of RNA product integrity using a 12-kb DNA template with modified nucleotides as substrates.

We next evaluated whether the integrity improvements were retained during transcription with modified nucleotides. N^1^-methylpseudouridine (m1Ψ) is an established modification in conventional therapeutic mRNA,^14^ whereas 5-methylcytidine (m5C) has emerged as a promising modification for saRNA, supporting protein expression while reducing innate immune activation in preclinical studies.^15^ We therefore tested M362K with m1ΨTP replacing UTP or m5CTP replacing CTP. Under both conditions, M362K produced transcripts with higher integrity than WT T7 RNAP (Fig. 3CD), demonstrating that the benefits of thumb-loop engineering were retained with these modified nucleotide substrates.

Together, these results establish thumb-loop engineering as a strategy to improve the integrity of long RNA transcripts, with several variants outperforming K389A on the 12K template and M362K showing the greatest benefit. Given the established roles of the thumb domain in EC stability and termination, proline substitutions may alter the loop conformational preferences and thereby reposition neighboring residues for productive interactions. M362K and K363R may tune the same interface by adding a positive charge or changing charged side chain geometry, respectively. These observations suggest that loop conformation and electrostatic interactions jointly influence transcriptional completion near the upstream DNA exit. The lack of an integrity reduction with K363A is compatible with compensatory contacts, although the proposed mechanism requires direct validation. M362K benefits both m1ΨTP and m5CTP, supporting its potential application to modified RNA synthesis. Further evaluation across diverse RNA sequences, together with measurements of EC kinetics and absolute full-length RNA yield, will clarify the mechanism, generality, and manufacturing value of this engineering strategy.

## Methods

### 1. Reagents

The relevant kits, including the dsRNA quantitative kit, site-directed mutagenesis kit, and BCA protein assay kit, as well as enzymes including RNase inhibitor (RI), inorganic pyrophosphatase (PPase), and nucleotides (A/G/C/UTP, 100 mM), were all provided by Vazyme Co., Ltd. (Nanjing, China). MgCl_2_, dithiothreitol (DTT), and other chemical reagents were purchased from Aladdin Industrial Corporation (Shanghai, China).

### 2. DNA template preparation

A self-replicating DNA template expressing the S protein was used as the modification model. The sequence includes the 5’ untranslated region (UTR), nonstructural protein 1-4 (nsp1-4) coding sequence, subgenomic promoter (SGP), S protein coding sequence, 3’ UTR, and poly(A) tail, totaling approximately 12,000 nt (SEQ ID NO.1, FigureS1). The sequence was divided by PCR as follows: (1) the complete template was sequentially divided from head to tail in 2,000-nt units, ultimately yielding six 2Kb templates named 2Kb-1 to 2Kb-6; (2) the complete template was sequentially divided from head to tail in 4,000-nt units, ultimately yielding three 4K templates named 4Kb-1 to 4Kb-3; (3) the first 6,000 nt of the complete template were taken to obtain a template named 6Kb; (4) the first 8,000 nt of the complete template were taken to obtain a template named 8Kb. Primer sequences are listed in Table S1.

### 3. In vitro mRNA preparation

Wild-type or variant T7 RNAP was added to the reaction system at 15 U/μL according to its enzymatic activity; the enzyme activity assay method was based on previous research.^16^ The reaction system contained final concentrations of 40 mM HEPES (pH 7.01), 2.5 mM tris(2-carboxyethyl)phosphine, 2 mM spermidine, 17.5 mM MgCl_2_, 50 ng/μL DNA template, 4 U/μL RI, 5 × 10^-3^ U/μL inorganic pyrophosphatase, 7.5 mM NTPs, and 5 mM cap analog. The reaction was carried out at 37 °C for 1.5 h. After the reaction, the obtained mRNA products were sent downstream for purification.

### 4. mRNA purification

After the mRNA transcription reaction, the product was diluted 10-fold with ddH_2_O and incubated at 37 °C for 5 min. VAHTS RNA Clean Beads were then added at 1.8× the product volume, mixed by pipetting, and allowed to stand for 5 min. The sample was placed on a magnetic rack for 5 min, and the supernatant was discarded. The beads were washed twice with 200 μL of 80% ethanol. After drying for 10 min, the product was removed from the magnetic rack, 50 μL of ddH_2_O was added, and the mixture was mixed by pipetting and allowed to stand for 5 min. The sample was then placed on a magnetic rack for 5 min, and the supernatant was collected to obtain the mRNA product. Finally, the concentration was measured using NanoDrop.

### 5. Determination of capping efficiency

The 1.25 μM mRNA product was mixed with probe at a 1:1 ratio and annealed. After annealing, 25 pmol of mRNA was digested with 20 U of RNase H at 25 °C for 20 min. For each 25 pmol of mRNA, 9 mL of streptavidin magnetic beads (TianDiRenHe) was used. First, 9 mL of beads was placed on a magnetic rack; after the solution became clear, the supernatant was discarded, and the beads were washed twice with 200 μL of ddH_2_O. The digested product was then mixed with the beads by pipetting and incubated with rotation for 30 min. Subsequently, the product was placed on a magnetic rack until the solution cleared, and the supernatant was discarded. The beads were washed twice with wash buffer (5 mM Tris-HCl, 0.5 mM EDTA, 1 M NaCl, pH 7.5) and twice with ddH_2_O. Finally, a corresponding amount of elution buffer (0.1 mM EDTA free acid, 1% methanol, ddH_2_O) was added to a final product concentration of 1.25 μM. The mixture was mixed by pipetting, reacted at 85 °C for 3 min, immediately placed on a magnetic rack, and the supernatant was collected. The supernatant was subjected to mass spectrometry analysis (mobile phase A: 0.065% HFIP + 0.0375% DIPEA in water; B: 0.065% HFIP + 0.0375% DIPEA in methanol; gradient: 0–20 min, B 5%→20%; column: Nano ChromCore C18, 50 × 4.6 mm, 1.8 μm; injection volume: 5 μL; flow rate: 0.3 mL/min; column temperature: 60 °C; m/z: 500–3000; instrument: Thermo Vanquish Flex (UPLC)-Orbitrap Exploris 120).

### 6. mRNA integrity detection

For mRNA with a fragment length ≤6000 nt, 100–200 ng of mRNA product was diluted to 5–10 ng/μL with dilution buffer (RNA Cartridge Kit #R1C405110, BiOptic Inc.) and denatured (65 °C for 5 min, followed by ice bath at 0 °C for 5 min). Integrity was then determined by capillary electrophoresis (injection 4 kV, 10 s; separation 4 kV, 480 s; 20 nt Marker).

For mRNA with a fragment length >6000 nt, the product was diluted to 50 ng/μL with ddH_2_O and denatured (65 °C for 5 min, followed by ice bath at 0 °C for 5 min). The sample was then diluted 11-fold with dilution buffer fragment (Analyzer RNA Kit DNF-471-0500, Agilent) and detected (Perform prerun: 8.0 kV, 30 s; Sample injection: 5.0 kV, 4 s; Dip count: 2; Separation: 8.0 kV, 40.0/50.0 min).

### 7. dsRNA detection

The dsRNA detection method was based on previously published research.^17^

## Author Contributions

Wei He: Methodology, Validation, Formal Analysis, Visualization, Writing – Original Draft. Hongli Zhou & Guiying Ji: Investigation. Xiaoyu Xu: Supervision, Funding Acquisition. Qiuheng Jin: Conceptualization, Project Administration, Supervision, Writing – Review & Editing.

