## Supplementary Figure and Table for "Engineering the thumb loop of T7 RNA polymerase improves the integrity of long RNA transcripts"

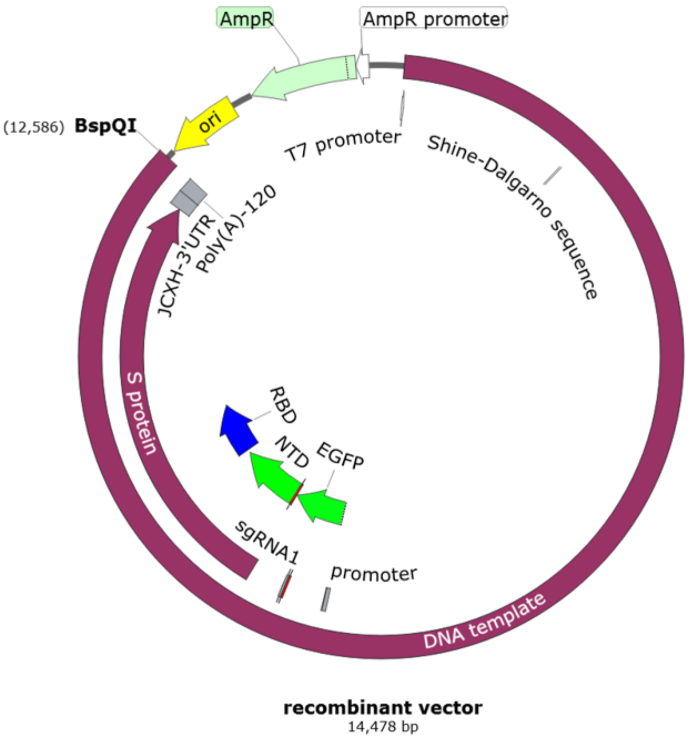


**Figure S1. Map of the plasmid used to generate linear template for 12K mRNA IVT.**

**Table S1. Primer sequences.**

| DNA | Primer | Sequence |
|---|---|---|
| 2K-1 | Forward | taatacgactcactataaggatggagaaagttcacgttgacatcgaggaagac |
| | Reverse | ttagtattcgccgtcgtgctcgctgg |
| 2K-2 | Forward | taatacgactcactataaggatgctgtacgacatcgacaggaaacagtgcg |
| | Reverse | taatatgagggtgcacatccggcttcg |
| 2K-3 | Forward | taatacgactcactataaggatggcagaggcttatgagtccatcgctaag |
| | Reverse | ttagagcgtgttctggatcgctgaaggc |
| 2K-4 | Forward | taatacgactcactataaggatgaacgtcctggcagctgccac |
| | Reverse | ttacttgtacagctcgtccatgccgagag |
| 2K-5 | Forward | taatacgactcactataaggatgttcgtgtttctggtgctgctgcctc |
| | Reverse | ttagccgatagggatgtcgcactcatagctattg |
| 2K-6 | Forward | taatacgactcactataagggccggcatctgtgcctcctacc |
| | Reverse | ttaggtgtagtgcagcttcacgcccttc |
| 4K-1 | Forward | taatacgactcactataaggatggagaaagttcacgttgacatcgaggaagac |
| | Reverse | taatatgagggtgcacatccggcttcg |
| 4K-2 | Forward | taatacgactcactataaggatggcagaggcttatgagtccatcgctaag |
| | Reverse | ttacttgtacagctcgtccatgccgagag |
| 4K-3 | Forward | taatacgactcactataaggatgttcgtgtttctggtgctgctgcctc |
| | Reverse | ttaggtgtagtgcagcttcacgcccttc |
| 6K | Forward | taatacgactcactataaggatggagaaagttcacgttgacatcgaggaagac |
| | Reverse | ttagagcgtgttctggatcgctgaaggc |
| 8K | Forward | taatacgactcactataaggatggagaaagttcacgttgacatcgaggaagac |
| | Reverse | ttacttgtacagctcgtccatgccgagag |
| 12K | Forward | taatacgactcactataaggatgggcggcgc |
| | Reverse | ctcacatgtgctcttctttttttt |

### DNA template sequence

tatgtaatacgactcactataaggatgggcggcgcatgagagaagcccagaccaattacctacccaaaatggaga
aagttcacgttgacatcgaggaagacagcccattcctcagagctttgcagcggagcttcccgcagtttgaggtagaa
gccaagcaggtcactgataatgaccatgctaatgccagagcgttttcgcatctggcttcaaaactgatcgaaacgga
ggtggacccatccgacacgatccttgacattggaagtgcgcccgcccgcagaatgtattctaagcacaagtatcatt
gtatctgtccgatgagatgtgcggaagatccggacagattgtataagtatgcaactaagctgaagaaaaactgtaag
gaaataactgataaggaattggacaagaaaatgaaggagctcgccgccgtcatgagcgaccctgacctggaaac
tgagactatgtgcctccacgacgacgagtcgtgtcgctacgaagggcaagtcgctgtttaccaggatgtatacgcgg
ttgacggaccgacaagtctctatcaccaagccaataagggagttagagtcgcctactggataggctttgacaccac
cccttttatgtttaagaacttggctggagcatatccatcatactctaccaactgggccgacgaaaccgtgttaacggctc
gtaacataggcctatgcagctctgacgttatggagcggtcacgtagagggatgtccattcttagaaagaagtatttgaa
accatccaacaatgttctattctctgttggctcgaccatctaccacgagaagagggacttactgaggagctggcacct
gccgtctgtatttcacttacgtggcaagcaaaattacacatgtcggtgtgagactatagttagttgcgacgggtacgtcg
ttaaaagaatagctatcagtccaggcctgtatgggaagccttcaggctatgctgctacgatgcaccgcgagggattct
tgtgctgcaaagtgacagacacattgaacggggagagggtctcttttcccgtgtgcacgtatgtgccagctacattgtg
tgaccaaatgactggcatactggcaacagatgtcagtgcggacgacgcgcaaaaactgctggttgggctcaacca
gcgtatagtcgtcaacggtcgcacccagagaaacaccaataccatgaaaaattaccttttgcccgtagtggcccag
gcatttgctaggtgggcaaaggaatataaggaagatcaagaagatgaaaggccactaggactacgagatagaca
gttagtcatggggtgttgttgggcttttagaaggcacaagataacatctatttataagcgcccggatacccaaaccatc
atcaaagtgaacagcgatttccactcattcgtgctgcccaggataggcagtaacacattggagatcgggctgagaa
caagaatcaggaaaatgttagaggagcacaaggagccgtcacctctcattaccgccgaggacgtacaagaagct
aagtgcgcagccgatgaggctaaggaggtgcgtgaagccgaggagttgcgcgcagctctaccacctttggcagct
gatgttgaggagcccactctggaagccgatgtcgacttgatgttacaagaggctggggccggctcagtggagacac
ctcgtggcttgataaaggttaccagctacgatggcgaggacaagatcggctcttacgctgtgctttctccgcaggctgt
actcaagagtgaaaaattatcttgcatccaccctctcgctgaacaagtcatagtgataacacactctggccgaaaag
ggcgttatgccgtggaaccataccatggtaaagtagtggtgccagagggacatgcaatacccgtccaggactttca
agctctgagtgaaagtgccaccattgtgtacaacgaacgtgagttcgtaaacaggtacctgcaccatattgccacac
atggaggagcgctgaacactgatgaagaatattacaaaactgtcaagcccagcgagcacgacggcgaatacctg
tacgacatcgacaggaaacagtgcgtcaagaaagaactagtcactgggctagggctcacaggcgagctggtgga
tcctcccttccatgaattcgcctacgagagtctgagaacacgaccagccgctccttaccaagtaccaaccataggg
gtgtatggcgtgccaggatcaggcaagtctggcatcattaaaagcgcagtcaccaaaaaagatctagtggtgagcg
ccaagaaagaaaactgtgcagaaattataagggacgtcaagaaaatgaaagggctggacgtcaatgccagaact
gtggactcagtgctcttgaatggatgcaaacaccccgtagagaccctgtatattgacgaagcttttgcttgtcatgcag
gtactctcagagcgctcatagccattataagacctaaaaaggcagtgctctgcggggatcccaaacagtgcggttttt
ttaacatgatgtgcctgaaagtgcattttaaccacgagatttgcacacaagtcttccacaaaagcatctctcgccgttg
cactaaatctgtgacttcggtcgtctcaaccttgttttacgacaaaaaaatgagaacgacgaatccgaaagagacta
agattgtgattgacactaccggcagtaccaaacctaagcaggacgatctcattctcacttgtttcagagggtgggtga
agcagttgcaaatagattacaaaggcaacgaaataatgacggcagctgcctctcaagggctgacccgtaaaggtg
tgtatgccgttcggtacaaggtgaatgaaaatcctctgtacgcacccacctcagaacatgtgaacgtcctactgaccc
gcacggaggaccgcatcgtgtggaaaacactagccggcgacccatggataaaaacactgactgccaagtaccct
gggaatttcactgccacgatagaggagtggcaagcagagcatgatgccatcatgaggcacatcttggagagaccg
gaccctaccgacgtcttccagaataaggcaaacgtgtgttgggccaaggctttagtgccggtgctgaagaccgctgg
catagacatgaccactgaacaatggaacactgtggattattttgaaacggacaaagctcactcagcagagatagtat
tgaaccaactatgcgtgaggttctttggactcgatctggactccggtctattttctgcacccactgttccgttatccattagg

aataatcactgggataactccccgtcgcctaacatgtacgggctgaataaagaagtggtccgtcagctctctcgcag
gtacccacaactgcctcgggcagttgccactggtcgtgtctatgacatgaacactggtacactgcgcaattatgatcc
gcgcataaacctagtacctgtaaacagaagactgcctcatgctttagtcctccaccataatgaacacccacagagtg
acttttcttcattcgtcagcaaattgaagggcagaactgtcctggtggtcggggaaaagttgtccgtcccaggcaaaat
ggttgactggttgtcagaccggcctgaggctaccttcagagctcggctggatttaggcatcccaggtgatgtgcccaa
atatgacataatatttgttaatgtgaggaccccatataaataccatcactatcagcagtgtgaagaccatgccattaag
cttagcatgttgaccaagaaagcttgtctgcatctgaatcccggcggaacctgtgtcagcataggttatggttacgctg
acagggccagcgaaagcatcattggtgctatagcgcggcagttcaagttttcccgggtatgcaaaccgaaatcctc
acttgaagagacggaagttctgtttgtattcattgggtacgatcgcaaggcccgtacgcacaatccttacaagctttcat
caaccttgaccaacatttatacaggttccagactccacgaagccggatgtgcaccctcatatcatgtggtgcgaggg
gatattgccacggccaccgaaggagtgattataaatgctgctaacagcaaaggacaacctggcggaggggtgtgc
ggagcgctgtataagaaattcccggaaagcttcgatttacagccgatcgaagtaggaaaagcgcgactggtcaaa
ggtgcagctaaacatatcattcatgccgtaggaccaaacttcaacaaagtttcggaggttgaaggtgacaaacagtt
ggcagaggcttatgagtccatcgctaagattgtcaacgataacaattacaagtcagtagcgattccactgttgtccac
cggcatcttttccgggaacaaagatcgactaacccaatcattgaaccatttgctgacagctttagacaccactgatgc
agatgtagccatatactgcagggacaagaaatgggaaatgactctcaaggaagcagtggctaggagagaagcag
tggaggagatatgcatatccgacgatagtagtgtgacagaacctgatgcggaactagtgagggtgcatccgaagag
ttctttggctggaaggaagggctacagcacaagcgatggcaaaactttctcatatttggaagggaccaagtttcacca
ggcggccaaggatatagcagaaattaatgccatgtggcccgttgcaacggaggccaatgagcaggtatgcatgtat
atcctcggagaaagcatgagcagtattaggtcgaaatgccccgtcgaagagtcggaagcctccacaccacctagc
acgctgccttgcttgtgcatccatgccatgactccagaaagagtacagcgcctaaaagcctcacgtccagaacaaa
ttactgtgtgctcatcctttccattgccgaagtatagaatcactggtgtgcagaagatccaatgctcccagcctatattgtt
ctcaccgaaagtgcctgcgtatattcatccaaggaagtatctcgtggaaacaccaccggtagacgagactccggag
ccatcggcagagaaccaatccacagaggggacacctgaacaaccaccacttataaccgaggatgagaccagg
actagaacgcctgagccgatcatcatcgaagaggaagaagaggatagcataagtttgctgtcagatggcccgacc
caccaggtgctgcaagtcgaggcagacattcacgggccgccctctgtatctagctcatcctggtccattcctcatgca
tccgactttgatgtggacagtttatccatacttgacaccctggagggagctagcgtgaccagcggggcaacgtcagc
cgagactaactcttacttcgcaaagagtatggagtttctggcgcgaccggtgcctgcgcctcgaacagtattcagga
accctccacatcccgctccgcgcacaagaacaccgtcacttgcacccagcagggcctgctcgagaaccagccta
gtttccaccccgccaggcgtgaatagggtgatcactagagaggagctcgaggcgcttaccccgtcacgcactccta
gcaggtcggtctcgagaaccagcctggtctccaacccgccaggcgtaaatagggtgattacaagagaggagtttg
aggcgttcgtagcacaacaacaatgacggtttgatgcgggtgcatacatcttttcctccgacaccggtcaagggcatt
tacaacaaaaatcagtaaggcaaacggtgctatccgaagtggtgttggagaggaccgaattggagatttcgtatgcc
ccgcgcctcgaccaagaaaaagaagaattactacgcaagaaattacagttaaatcccacacctgctaacagaag
cagataccagtccaggaaggtggagaacatgaaagccataacagctagacgtattctgcaaggcctagggcatta
tttgaaggcagaaggaaaagtggagtgctaccgaaccctgcatcctgttcctttgtattcatctagtgtgaaccgtgcct
tttcaagccccaaggtcgcagtggaagcctgtaacgccatgttgaaagagaactttccgactgtggcttcttactgtatt
attccagagtacgatgcctatttggacatggttgacggagcttcatgctgcttagacactgccagtttttgccctgcaaa
gctgcgcagctttccaaagaaacactcctatttggaacccacaatacgatcggcagtgccttcagcgatccagaac
acgctccagaacgtcctggcagctgccacaaaaagaaattgcaatgtcacgcaaatgagagaattgcccgtattg
gattcggcggcctttaatgtggaatgcttcaagaaatatgcgtgtaataatgaatattgggaaacgtttaaagaaaacc
ccatcaggcttactgaagaaaacgtggtaaattacattaccaaattaaaaggaccaaaagctgctgctctttttgcga
agacacataatttgaatatgttgcaggacataccaatggacaggtttgtaatggacttaaagagagacgtgaaagtg
actccaggaacaaaacatactgaagaacggcccaaggtacaggtgatccaggctgccgatccgctagcaacag

cgtatctgtgcggaatccaccgagagctggttaggagattaaatgcggtcctgcttccgaacattcatacactgtttgat
atgtcggctgaagactttgacgctattatagccgagcacttccagcctggggattgtgttctggaaactgacatcgcgt
cgtttgataaaagtgaggacgacgccatggctctgaccgcgttaatgattctggaagacttaggtgtggacgcagag
ctgttgacgctgattgaggcggctttcggcgaaatttcatcaatacatttgcccactaaaactaaatttaaattcggagc
catgatgaaatctggaatgttcctcacactgtttgtgaacacagtcattaacattgtaatcgcaagcagagtgttgagag
aacggctaaccggatcaccatgtgcagcattcattggagatgacaatatcgtgaaaggagtcaaatcggacaaatt
aatggcagacaggtgcgccacctggttgaatatggaagtcaagattatagatgctgtggtgggcgagaaagcgcctt
atttctgtggagggtttattttgtgtgactccgtgaccggcacagcgtgccgtgtggcagacccccctaaaaaggctgttt
aagcttggcaaacctctggcagcagacgatgaacatgatgatgacaggagaagggcattgcatgaggaatcaac
acgctggaaccgagtgggtattctttcagagctgtgcaaggcagtagaatcaaggtatgaaaccgtaggaacttcca
tcatagttatggccatgactactctagctagcagtgttaaatcattcagctacctgagaggggcccctataactctctac
ggctaacctgaatggactacgacatagtctagtccgccaaggccaccatggtgagcaagggcgaggagctgttca
ccggggtggtgcccatcctggtcgagctggacggcgacgtaaacggccacaagttcagcgtgtccggcgagggc
gagggcgatgccacctacggcaagctgaccctgaagttcatctgcaccaccggcaagctgcccgtgccctggcc
caccctcgtgaccaccctgacctacggcgtgcagtgcttcagccgctaccccgaccacatgaagcagcacgactt
cttcaagtccgccatgcccgaaggctacgtccaggagcgcaccatcttcttcaaggacgacggcaactacaagac
ccgcgccgaggtgaagttcgagggcgacaccctggtgaaccgcatcgagctgaagggcatcgacttcaaggagg
acggcaacatcctggggcacaagctggagtacaactacaacagccacaacgtctatatcatggccgacaagcag
aagaacggcatcaaggtgaacttcaagatccgccacaacatcgaggacggcagcgtgcagctcgccgaccact
accagcagaacacccccatcggcgacggccccgtgctgctgcccgacaaccactacctgagcacccagtccgc
cctgagcaaagaccccaacgagaagcgcgatcacatggtcctgctggagttcgtgaccgccgccgggatcactct
cggcatggacgagctgtacaagatgttcgtgtttctggtgctgctgcctctggtgagctcccagtgcgtgaatctgacca
caaggacccagctgccccctgcctataccaactccttcacacggggcgtgtactatcccgacaaggtgttccggag
cagcgtgctgcactccacacaggatctgtttctgcctttcttttctaacgtgacctggttccacgccatccacgtgagcg
gcaccaatggcacaaagaggttcgacaacccagtgctgcccttcaatgatggcgtgtacttcgcctccaccgagaa
gtctaatatcatccgcggctggatctttggcaccacactggacagcaagacacagtccctgctgatcgtgaacaatg
ccaccaacgtggtcatcaaggtgtgcgagttccagttttgtaacgatccattcctgggcgtgtactatcacaagaaca
ataagtcttggatggagagcgagtttcgcgtgtattcctctgccaacaattgcacatttgagtacgtgtcccagcccttcc
tgatggacctggagggcaagcagggcaatttcaagaacctgcgggagttcgtgtttaagaacatcgatggctacttc
aaaatctactccaagcacaccccaatcaatctggtgagagacctgccacagggcttctctgccctggagccactgg
tggatctgcccatcggcatcaacatcacccggtttcagacactgctggccctgcacagaagctacctgacaccagg
cgacagctcctctggatggaccgcaggagcagcagcctactatgtgggctatctgcagcccaggaccttcctgctg
aagtacaacgagaatggcaccatcacagacgcagtggattgcgcactggacccccctgtctgagaccaagtgtac
actgaagtcctttaccgtggagaagggcatctatcagacaagcaacttcagggtgcagcctaccgagtccatcgtgc
gctttcccaatatcacaaacctgtgcccttttggcgaggtgttcaatgcaaccaggttcgcaagcgtgtacgcatggaa
taggaagcgcatctccaactgcgtggccgactattctgtgctgtacaatagcgcctccttctctacctttaagtgctacg
gcgtgagccccacaaagctgaatgacctgtgctttaccaacgtgtacgccgattccttcgtgatcaggggcgacgag
gtgcgccagatcgcaccaggacagacaggcaagatcgcagactacaactataagctgcctgacgatttcaccgg
ctgcgtgatcgcctggaacagcaacaatctggatagcaaagtgggcggcaactacaattatctgtaccggctgttta
gaaagtctaacctgaagccattcgagagggacatctccacagaaatctaccaggccggctctacccccctgcaatg
gcgtggagggctttaactgttatttccctctgcagagctacggcttccagccaacaaatggcgtgggctatcagcccta
ccgcgtggtggtgctgtcttttgagctgctgcacgcccctgcaacagtgtgcggaccaaagaagtccaccaatctggt
gaagaacaagtgcgtgaacttcaacttcaacggactgaccggcacaggcgtgctgaccgagtccaacaagaagt
tcctgccttttcagcagttcggcagggacatcgcagataccacagacgccgtgcgcgaccctcagaccctggagat

cctggatatcacaccatgctccttcggcggcgtgtctgtgatcacaccaggcaccaatacaagcaaccaggtggcc
gtgctgtatcaggacgtgaactgtaccgaggtgcccgtggcaatccacgcagatcagctgacccctacatggcggg
tgtactctaccggcagcaacgtgttccagacaagagccggatgcctgatcggagcagagcacgtgaacaatagct
atgagtgcgacatccctatcggcgccggcatctgtgcctcctaccagacccagacaaactccccaaggagagca
cggtctgtggcaagccagtccatcatcgcctataccatgagcctgggcgccgagaactccgtggcctactccaaca
attctatcgccatccctaccaatttcacaatctccgtgaccacagagatcctgccagtgagcatgaccaagacatcc
gtggactgcacaatgtatatctgtggcgattccaccgagtgctctaatctgctgctgcagtacgggtccttttgtacccag
ctgaacagagccctgacaggcatcgccgtggagcaggacaagaatacacaggaggtgttcgcccaggtgaagc
aaatctacaagaccccacccatcaaggactttggcggcttcaactttagccagatcctgcccgatcctagcaagcc
atccaagcggtcttttatcgaggacctgctgttcaataaggtgaccctggccgatgccggcttcatcaagcagtatggc
gattgcctgggcgacatcgccgccagagacctgatctgtgcccagaagtttaacggcctgaccgtgctgcctccact
gctgacagatgagatgatcgcccagtacacatctgccctgctggcaggcaccatcacaagcggatggaccttcgg
cgcaggagccgccctgcagatcccctttgccatgcagatggcctatcggttcaatggcatcggcgtgacccagaat
gtgctgtacgagaaccagaagctgatcgccaatcagtttaactccgccatcggcaagatccaggactctctgagctc
cacagcaagcgccctgggcaagctgcaggatgtggtgaatcagaacgcccaggccctgaacaccctggtgaag
cagctgtctagcaatttcggcgccatctcctctgtgctgaacgatatcctgagccggctggacaaggtggaggcaga
ggtgcagatcgaccggctgatcacaggcagactgcagtccctgcagacctacgtgacacagcagctgatcaggg
cagcagagatcagggcatctgccaacctggcagcaaccaagatgagcgagtgcgtgctgggccagtccaagag
agtggacttttgtggcaagggctatcacctgatgagcttcccacagtccgcccctcacggagtggtgtttctgcacgtg
acctacgtgccagcccaggagaagaacttcaccacagcaccagcaatctgccacgatggcaaggcacactttcc
tagggagggcgtgttcgtgagcaatggcacccactggtttgtgacacagcgcaacttctacgagccacagatcatca
ccacagacaatacattcgtgtccggcaactgtgacgtggtcatcggcatcgtgaacaataccgtgtatgatcctctgc
agccagagctggactcttttaaggaggagctggataagtacttcaagaaccacaccagccccgacgtggatctgg
gcgacatctctggcatcaatgccagcgtggtgaacatccagaaggagatcgacaggctgaatgaggtggccaag
aatctgaacgagtccctgatcgatctgcaggagctgggcaagtatgagcagtacatcaagtggccctggtatatctg
gctgggcttcatcgccggcctgatcgccatcgtgatggtgaccatcatgctgtgctgtatgacaagctgctgttcctgcc
tgaagggctgctgttcttgtggcagctgctgtaagtttgatgaggacgatagcgagcctgtgctgaagggcgtgaagct
gcactacacctaaggcgcgcccacccagcggccgcatacagcagcaattggcaagctgcttacatagaactcgc
ggcgattggcatgccgccttaaaatttttattttatttttcttttcttttccgaatcggattttgtttttaatatttcaaaaaaaaaaa
aaaaaaaaaaaaaaaaaaaaagcatatgactaaaaaaaaaaaaaaaaaaaaaaaaaaaaaaaaaaaaaaaaaa
aaaaaaaaaaaaaaaaaaaaaaaaaaaaaaaaaaaaaaaaaaaaaaaaaaaaaaaaa
